# Task-related changes in aperiodic activity are related to visual working memory capacity independent of event-related potentials and alpha oscillations

**DOI:** 10.1101/2022.01.18.476852

**Authors:** Sian Virtue-Griffiths, Alex Fornito, Sarah Thompson, Mana Biabani, Jeggan Tiego, Tribikram Thapa, Neil W. Bailey, Nigel C. Rogasch

**Affiliations:** School of Psychological Sciences, Turner Institute of Brain and Mental Health, and Monash Biomedical Imaging, Monash University, VIC, Australia; Clinical Sciences, Murdoch Children’s Research Institute, Melbourne, VIC, Australia; School of Medicine and Psychology, The Australian National University, Canberra, ACT; Monarch Research Institute, Monarch Mental Health Group, Sydney, NSW; School of Biomedicine, University of Adelaide, SA, Australia; Hopwood Centre for Neurobiology, Lifelong Health Theme, South Australian Health and Medical Research Institute, SA, Australia

**Keywords:** Electroencephalography (EEG), Aperiodic activity, Working memory, Alpha oscillations, Event-related potentials

## Abstract

Research using electroencephalography (EEG) has shown that individual differences in visual working memory capacity are related to slow-wave event-related potentials (ERPs) and suppression of alpha-band oscillatory power during the delay period of memory tasks. However, recent evidence suggests that changes in non-oscillatory (aperiodic) features of the EEG signal are related to working memory performance. We assessed several features of task-related changes in aperiodic activity including its spatial distribution, the effect of memory load, and the relationships between aperiodic activity, memory capacity, slow-wave ERPs, and alpha suppression. Eighty-four healthy individuals performed a continuous recall working memory (WM) task consisting of 2, 4 or 6 coloured squares while EEG was recorded. Aperiodic activity during a baseline and WM delay period was quantified by fitting a model to the background of the EEG power spectra, which returned parameters describing the slope (exponent) and broadband offset of the spectra. The aperiodic exponent decreased (i.e., slope flattened) in lateral parieto-occipital electrodes but increased (i.e., slope steepened) in fronto-central electrodes during the WM delay period, whereas the offset decreased over parieto-occipital electrodes. These task-related changes in aperiodic activity did not differ between memory loads. Larger increases in the aperiodic exponent were associated with higher working memory capacity measured from both the WM task and a separate battery of complex span tasks, a relationship that was independent of slow-wave ERPs and alpha suppression. Our findings suggest that WM task-related changes in aperiodic activity are region specific and reflect an independent neural mechanism that is important for general working memory ability.

## 1. Introduction

Working memory refers to the active maintenance of information in ‘the mind’ for brief periods to serve an ongoing task (Luck and Vogel 2013). While working memory is considered domain general, there is also evidence for modality specific domains, for example, a “phonological loop” for verbal information and “visuospatial sketchpad” for visual information (Baddeley 2012). Human working memory is widely accepted to have a limited capacity. The maximum amount of information that can be held at one time in visual working memory (VWM), known as working memory capacity, is limited between 1 to 5 items, with the value varying between individuals (Logie 2011). Differences in VWM capacity appear to be important for broader cognitive ability, accounting for approximately 45% of the variance in fluid intelligence (Fukuda et al. 2010). Furthermore, VWM capacity is reduced with healthy aging and in many neurological and psychiatric disorders (Berryman et al. 2013; Ding et al. 2015; Park and Gooding 2014; Rock et al. 2014). These reductions are often associated with reduced functional outcomes, such as difficulties maintaining work and social connections (Green 2006). Taken together, these findings suggest VWM is an essential cognitive ability for daily living.

Studies using electroencephalography (EEG) to record brain activity have identified several neural correlates of visual working memory capacity. For example, the maintenance of items in working memory is accompanied by a slow-wave event-related potential (ERP) during the working memory delay period (the period between the presentation of items to remember and the recognition probe or recall period), which is negative in polarity over posterior electrodes (Ruchkin et al. 1992; 1995). This slow-wave ERP lateralises to the contralateral hemisphere when items are presented to a given visual field (often called contralateral delay activity) (Vogel and Machizawa 2004; Vogel, McCollough, and Machizawa 2005; McCollough, Machizawa, and Vogel 2007) and becomes more negative as memory load increases, until an individual’s capacity is exceeded (Vogel, McCollough, and Machizawa 2005). The slow-wave ERP also correlates with individual working memory capacity (Vogel and Machizawa 2004; Vogel, McCollough, and Machizawa 2005; Fukuda, Mance, and Vogel 2015). Slow-wave ERPs recorded during working memory tasks are also correlated with measures indexing attentional control, long-term memory, and fluid intelligence, indicating that individual differences in delay activity are important for a broad range of cognitive abilities (Unsworth et al. 2015).

In addition to slow wave ERPs, working memory of visual items is accompanied by a suppression of oscillations in the alpha frequency range (8-12 Hz) during the encoding and early delay period (Gevins et al. 1997; Sauseng et al. 2009). Like slow wave ERPs, alpha suppression lateralises to the contralateral hemisphere when items are presented to a cued visual field. Alpha suppression also increases with memory load until an individual’s capacity is exceeded, and correlates with working memory capacity, with greater suppression being associated with greater capacity (Fukuda, Mance, and Vogel 2015). Increases in alpha-band oscillatory power are also observed over regions ipsilateral to relevant information in lateralized tasks, and in the delay period in some working memory tasks, suggesting changes in alpha-band power may reflect a mechanism for gating information in working memory (Jensen and Mazaheri 2010). The similarities between slow-wave ERPs and alpha suppression have led to the proposal that these measures may reflect a shared neural mechanism (van Dijk et al. 2010). However, the slow wave ERP and alpha suppression account for independent variance in working memory capacity and occur across different time spans (Fukuda, Mance, and Vogel 2015). Furthermore, recent studies using multivariate pattern analysis indicate that an item’s spatial location can be decoded from alpha activity (Bae and Luck 2018; Foster et al. 2016), whereas both spatial location and stimulus features (e.g., orientation) can be decoded from slow-wave ERPs (Bae and Luck 2018), suggesting the two measures reflect partially overlapping, but independent neural mechanisms supporting working memory.

Although most EEG studies of working memory have focused on ERPs and/or oscillatory activity, neural signals also contain non-oscillatory, aperiodic activity. This activity contains voltage shifts with phase angles that are identified by traditional frequency band-power analyses as reflecting power at specific frequencies, but instead of oscillating with a regular period, the voltage shifts do not oscillate with a regular period. In the frequency domain, this aperiodic activity is represented by a 1/f-like distribution, with power decreasing exponentially as frequencies increase. Importantly, aperiodic activity can change over both short timescales (e.g., during tasks (Podvalny et al. 2015)) and long timescales (e.g., over the human lifespan (Voytek et al. 2015; Merkin et al. 2023)). Several methods have been proposed to separate aperiodic parameters from periodic oscillatory activity in neural power spectra, including linear regression approaches (Voytek et al. 2015; Gao, Peterson, and Voytek 2017), IRASA (Irregular Resampling Auto-Spectral Analysis) (Wen and Liu 2016), SpecParam/FOOOF (Fitting Oscillations & One-Over-F) (Donoghue et al. 2020), and PaWNextra (Pink and White Noise Extractor) (Barry and Blasio 2021). Using one approach, Voytek et al. (2015) showed the aperiodic exponent (representing the steepness of the 1/f-like slope in log-power-log-frequency space) was associated with age-related changes in working memory performance. In a more recent study, Donoghue et al. (2020) showed a decrease in both the aperiodic exponent (suggesting a flattening of the 1/f-like slope) and a decrease in the broadband offset (suggesting an overall reduction in broadband power) in posterior electrodes contralateral to attended items during the delay period of a working memory task compared to a baseline period. Importantly, even when changes in alpha-band oscillatory activity were included in the model, the changes in aperiodic activity were independently associated with working memory performance. Taken together, these studies suggest that changes in aperiodic activity may be important for working memory performance. However, most studies have focused on posterior electrodes, and it remains unclear how aperiodic activity is modulated across the scalp during working memory. For instance, the broadband offset at rest is correlated with measures of cognitive control in frontocentral electrodes, suggesting other sites across the scalp may be relevant for cognitive tasks (Clements et al. 2021). Furthermore, it is unclear whether aperiodic activity is sensitive to variation in working memory load, and whether changes in aperiodic activity share a common neural mechanism with other EEG measures such as slow-wave ERPs.

The aim of this study was to comprehensively assess how aperiodic activity is modulated during a visual working memory task and how aperiodic activity relates to working memory performance. We assessed four main questions: 1) How does aperiodic activity change across the scalp during the delay period of a working memory task compared to a baseline period? 2) Are changes in aperiodic activity from baseline to the working memory delay period modulated by increasing working memory load? 3) Are changes in aperiodic activity from baseline to the working memory delay period associated with working memory performance across individuals and tasks? and 4) Are changes in aperiodic activity from baseline to the working memory delay period independent from other EEG measures such as slow-wave ERPs and alpha suppression?

## 2. Methods

### 2.1 Participants

Eighty-four right-handed participants (28 males) aged 19-40 (*M_age_* = 26.7, *SD* ± 4.9) were recruited for a multi-modal experiment including EEG, magnetic resonance imaging (MRI), transcranial magnetic stimulation (TMS), cognitive testing, and genetics. Only EEG and cognitive data were analysed in the current study. Exclusion criteria included: a history of neurological or psychiatric illness; colour-blindness; and contraindications to either MRI (Sawyer-Glover and Shellock 2000), or TMS (Rossi et al. 2009). All participants had a predominantly Caucasian background (four grandparents of European descent) as the broader recruitment pool that participants were recruited from was also used for a separate genetics study. Experimental procedures were approved by the Monash University Human Research Ethics Committee (MURHEC number 8836).

### 2.2 Experimental design

In the EEG session, participants were seated in a chair positioned approximately 480 mm from a computer monitor (monitor dimensions = 540 x 640 mm). Following preparation of the electrodes, EEG was recorded for an 8-minute “rest” period where participants sat quietly with their eyes open (data not analysed). Following the rest condition, participants performed a continuous recall working memory task (described below). First, participants were provided with instructions on how to complete the task. Next, participants completed 2 control tasks testing motor precision (20 trials) and colour perception (30 trials) respectively, followed by a brief (∼1 min) practice version of the continuous recall working memory task. Participants then completed four blocks of the continuous recall task while EEG was recorded, taking 20-25 minutes to complete. In a separate cognitive assessment session held at least one day before the EEG session, participants completed a battery of computerised cognitive tasks including 3 complex span tasks which were also used to index working memory ability (see description below).

### 2.3 EEG acquisition

EEG was recorded from 62 passive electrodes in standard 10-10 positions (c-ring slit electrodes, EASYCAP, Germany). The electrodes were referenced to FPz and grounded to AFz. The EEG signal was amplified (1000×), filtered (DC-2000 Hz), and digitised (10 kHz; Syamps2, Compumedics Ltd.) for offline analysis. Electrode impedance was reduced to <5 kΩ by applying abrasive gel with cotton tips and then conductive gel with blunt-ended syringes between the electrode and the skin.

### 2.4 Continuous recall working memory task

Participants performed a whole-field, partial report, continuous recall working memory task for colour (fig. 1; adapted from the MemToolbox; https://visionlab.github.io/MemToolbox/) (Suchow et al. 2013). Each trial began with a central fixation cross presented for 1500 ms (baseline period). Following the baseline period, a memory array of either 2, 4, or 6 coloured squares was presented for 500 ms (encoding period). The colours were randomly drawn from a continuous colour array (360 individual colours equally spaced across the colour spectrum; minimum difference of 24° between colours presented for each trial) and were presented in one of eight positions equally spaced around the fixation cross. Participants were instructed to remember as many squares as possible over the delay period, which consisted of a blank screen presented for 1000 ms. Following the delay period, greyed-out squares were presented in identical positions to the memory array with one target square highlighted by a black outline. Participants were asked to indicate the colour of the highlighted square from the memory array by moving a mouse cursor over a colour wheel with their right hand (note the orientation of the colour wheel was changed randomly between trials). The colour of the highlighted square updated instantaneously to match the position of the cursor on the colour wheel as the participant moved the cursor. Participants finalised their response by pressing the left mouse button and the baseline period for the next trial began. One hundred trials for each load were presented (total = 300 trials). Trials were presented in four blocks of 75 trials (25 trials of each load in a random order), with approximately 2-3 mins rest between each block.

**Figure 1.**
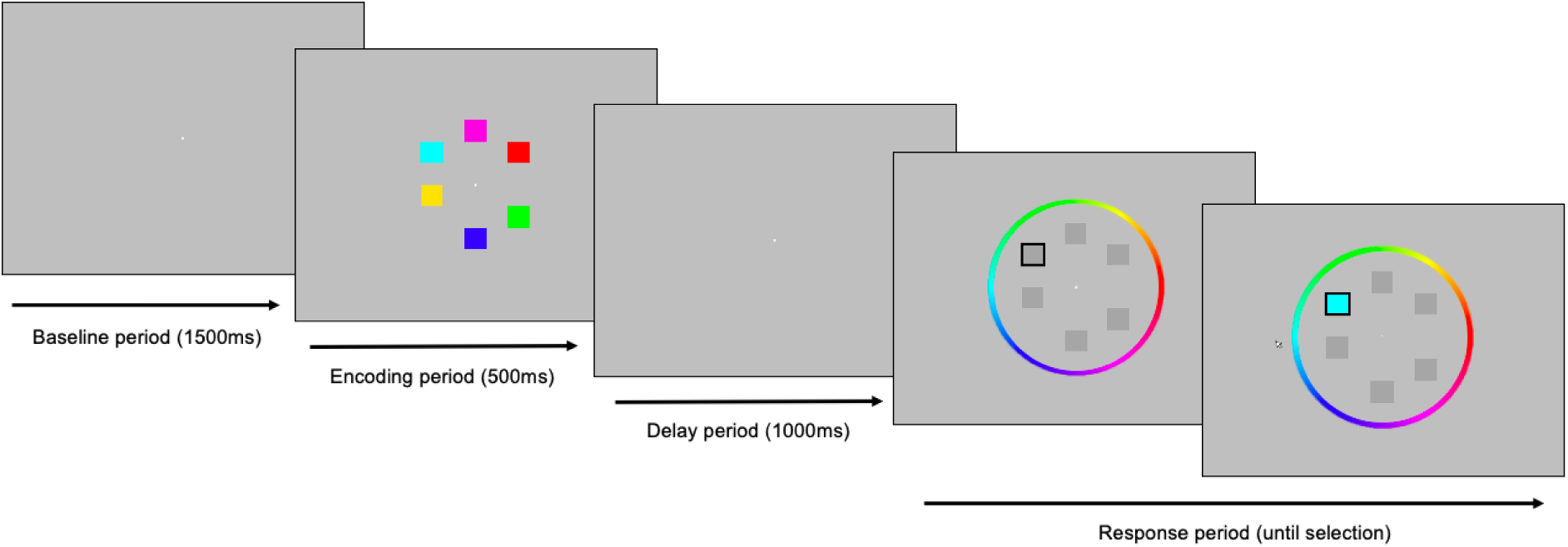
Continuous recall working memory task. Following a baseline period of 1500 ms, an array of either 2, 4 or 6 coloured squares was presented for 500 ms (encoding period). The squares were then removed for 1000 ms (delay period), after which a target square was indicated (black outline) and participants were instructed to match the colour as closely as possible on a colour wheel by navigating a mouse cursor (response period). The colour of the target square was instantaneously updated as the participant navigated around the colour wheel until the response was finalised by a button press.

Performance on each trial was quantified by calculating the distance between the target and the selected colour in degrees, resulting in a distribution of performance scores across trials for each load. Two measures of overall working memory ability (capacity and precision) were calculated for each load by fitting a standard mixture model (Zhang and Luck 2008) to the performance distribution using a maximum likelihood approach implemented in the MemToolbox (Suchow et al. 2013). This approach returns two parameters optimized to the data: the guess rate, *g*, which reflects the probability the participant remembers nothing about the items and guesses randomly, and the standard deviation, *σ*, which is inversely proportional to the precision of the participants memory. Capacity (*K*) was calculated by multiplying the probability the participant had a representation of the item in memory (1-*g*) by the set size for each load (2, 4 or 6). Following the approach of Fukuda et al (2015), we also calculated the average *K* for supracapacity set sizes (loads 4 and 6) to provide a single measure of visual working memory capacity for each individual (deemed *Kcd*).

### 2.5 EEG processing

EEG data were analysed using the EEGLAB (Delorme and Makeig 2004), the EEG cleaning-relevant tools from TESA (Rogasch et al. 2017), and FieldTrip (Oostenveld et al. 2011) toolboxes in MATLAB (R2015b, The Mathworks, USA) and the SpecParam/FOOOF toolbox (Donoghue et al. 2020) in Python (v3.6.5, Python Software Foundation). First, EEG data were downsampled to 1000 Hz. The data were then band-pass (0.1-100 Hz) and band-stop (48-52 Hz) filtered using the *eegfiltnew.m* function. After filtering, data were epoched around the beginning of the encoding period (−1000 to 2000 ms; which includes the baseline, encoding, delay, and first 500 ms of the response period), and baseline corrected by subtracting the mean baseline amplitude from the active period for each individual electrode (−500 to 0 ms). Data from the four separate working memory task blocks were then concatenated into a single file. Bad channels (electrodes with poor skin contact or that became disconnected) and bad trials (epochs contaminated by excessive muscle activity, large blinks/eye movements, motion artifacts etc.) were removed from the data following visual inspection. Residual artifacts were further minimised by the identification and subtraction of artifact components using independent component analysis (FastICA) (Hyvarinen 1999). Independent components were automatically categorised into neural and artifact components based on temporal, spatial and spectral characteristics using heuristic rules implemented in TESA (Rogasch et al. 2017). Categorisation of artifactual components was then visually assessed and components representing persistent muscle activity, eye blinks, lateral eye movements, or electrode noise were removed from the unmixing matrix before the scalp space data were reconstructed. Of note, persistent muscle activity can substantially alter the EEG power spectra and components representing muscle activity were selected for removal using an approach validated using pharmacologic paralysis (Fitzgibbon et al. 2016). Finally, missing electrodes were replaced using spherical interpolation, and the data were re-referenced to the average of all electrodes. Remaining trials were then separated into each working memory load condition (2, 4 and 6).

### 2.6 Aperiodic activity and alpha-band oscillatory power

To characterise task-related changes in the EEG power spectra including aperiodic and oscillatory activity, EEG data from each epoch were converted into the frequency domain using Welch’s method (*pwelch.m* function in MATLAB) across two 800 ms time periods: a baseline period (−800 to 0 ms relative to the onset of the encoding period stimuli) and a delay period (650 to 1450 ms after the onset of the encoding period stimuli, matching the working memory delay period). The delay period timing was chosen to capture the EEG signals of interest (slow-wave ERP, alpha suppression) while minimising ERPs related to encoding. The resulting power spectral density (PSD) estimates were then averaged over trials for each condition. To quantify aperiodic activity, the 1/f-like broadband background of the PSD from each time period and each electrode was fit between 3-30 Hz using the FOOOF toolbox. We limited the frequency range to >3 Hz to ensure at least 2 cycles of the minimum oscillation frequency of interest within the analysis window (Roach and Mathalon 2008) and <30 Hz to further minimise the contribution of any residual muscle activity on the spectra (Voytek et al. 2015). The methods used in the SpecParam/FOOOF toolbox are specifically designed to exclude oscillatory peaks from the fit, thereby ensuring an accurate estimate of aperiodic activity (Donoghue et al. 2020). The fitting approach returns two parameters: an exponent value which quantifies the steepness of the 1/f-like slope in log-log space; and an offset value which quantifies the broadband offset of the spectra. To quantify alpha-band oscillatory activity, mean power between 8-12 Hz was calculated from the PSD for each time period and each electrode.

### 2.7 Slow wave ERPs

Slow wave ERPs were quantified by taking the mean EEG signal across trials for each electrode (i.e., the ERP) and then calculating the mean EEG voltage between 650 to 1450 ms, corresponding to the delay period.

### 2.8 Complex span working memory tasks

Three complex span working memory tasks (backward digit span, symmetry span, and operation span; fig. 2) were also administered in a separate session and used to determine whether relationships between EEG measures and working memory performance generalised beyond the continuous recall task to broader measures of working memory ability. We used these tasks as capacity measures from complex span tasks and continuous recall/change detection tasks (also called visual array tasks) are thought to measure related components of working memory ability (Shipstead et al. 2012; Shipstead, Harrison, and Engle 2015). Confirmatory factor analysis conducted in Mplus 8.3 (Muthén and Muthén 2002) was used to quantify complex span performance, which extracts shared variance from related tasks, thus minimising the influence of task-specific components on performance estimates (Bollen and Noble 2011). As such, confirmatory factor analysis was used to obtain a single estimate of complex span performance for each individual, which represented a composite of the shared variance from each of these non-EEG tasks. To ensure adequate power to perform confirmatory factor analysis, we included data from an additional 34 participants who also completed the complex span tasks (total n = 118), resulting in an adequate case-to-parameter ratio of 39:1 (Jackson 2001). Thus, complex span performance for a given individual represented a normalised value in relation to the overall sample included. Complex span tasks were completed using the Inquisit lab test library (Inquisit 5.011, 2016). Three tasks contributed to participants complex span factor score estimates: Backwards Digit Span (Woods et al. 2011), Operation Span (Unsworth et al. 2005) and Symmetry Span (Kane et al. 2004). Performance on each task was calculated by summing the load values of all accurately recalled trials. Factor score estimates were generated from the confirmatory factor analysis of the complex span factor using the regression method (DiStefano and Motl 2009). Factor score determinacy was reasonably high (ρ = 0.816), indicating that the factor score estimates corresponded closely with the underlying latent variable (Grice 2001).

**Figure 2.**
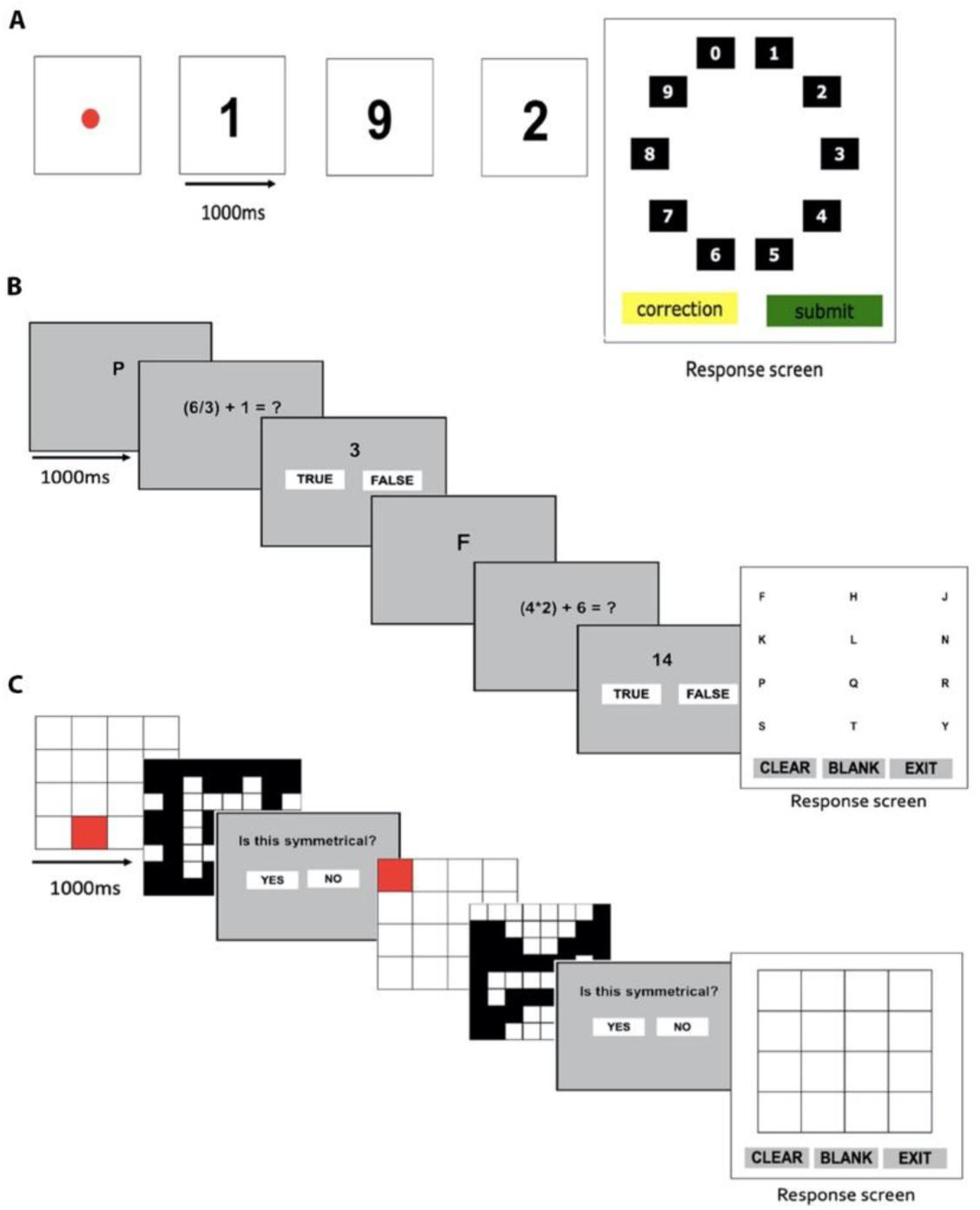
Complex span working memory tasks. Example trial of backwards digit span task (A), operation span task (B), and symmetry span task (C).

#### 2.8.1 Backwards Digit Span task

Participants were instructed to remember a sequence of single digit numbers, and then recall the numbers in reverse order by selecting them from a test screen (fig. 2A). Fourteen trials were conducted with between 2 to 8 numbers presented in each trial (i.e., 2 trials at each load).

#### 2.8.2 Operation span task

Participants were required to remember the order of a series of alphabetical letter. Between letters, equations were presented and participants were required to judge whether the provided answer was true or false (fig. 2B). Following the final equation, participants were required to recall the letters in correct order from a test screen. Fifteen trials were conducted with between 3 to 7 letters in each trial (i.e., 3 trials at each load).

#### 2.8.3 Symmetry Span task

Participants were required to remember the order and location of a series of highlighted square on a 4×4 grid. Between each presented square, participants were required to judge whether another grid pattern was symmetrical or asymmetrical across the vertical axis (fig.2C). Following the final symmetry judgement, participants were required to indicate the location of the presented squared in order from a test screen. Twelve trials were conducted with between 2 to 5 squares presented in each trial (i.e., 3 trials at each load).

### 2.9 Statistics

To assess the effect of increasing memory load on working memory performance in the continuous recall task, a one-way repeated measures analysis of variance (ANOVA) was conducted on capacity and precision measurements (MATLAB R2024b). For EEG analysis, non-parametric, cluster-based permutation statistics were performed using Fieldtrip to determine the effect of task and load on aperiodic exponent, offset, alpha-band power and slow wave ERPs (Oostenveld et al. 2011). Clusters were defined as two or more neighbouring electrodes in which the t-statistic exceeded a threshold of p<0.05 (two-sided dependent t-test). Monte Carlo p-values were calculated on 5000 random permutations and a value of p<0.05 (two-tailed unless indicated otherwise) was used as a cluster significance threshold for all analyses and adjusted to account for multiple tests (e.g., if identical tests were performed over 3 loads, p<0.017 was considered significant [0.05/3=0.017]). To quantify effect sizes, mean Cohen’s d values were calculated across all electrodes contributing to a significant cluster. To assess the relationship between EEG measures and working memory performance (working memory capacity, *Kcd*; complex span factor loading), we used the cluster-based permutation approach as outlined above. Electrode clusters were defined using a Pearson’s correlation with a threshold of p<0.05 adjusted for multiple tests. Participants with 4 or more electrodes with a z-score >3 for a given measure of interest (e.g., slow-wave ERP, 1/f slope, alpha power) were removed to minimise the impact of extreme outliers on statistical analyses. If a relationship was observed with working memory capacity, we used one-tailed tests for subsequent comparisons with complex span performance given the hypothesised direction of the relationship was known. Mean Pearson’s r values were calculated across all electrodes contributing to a significant cluster. Finally, a multiple linear regression analysis was performed to assess whether aperiodic activity, alpha suppression and slow-wave ERPs were independently related to working memory performance. Input variables for each EEG measure were calculated using two different approaches: either by averaging across all electrodes contributing to significant clusters from the Pearson’s correlation analysis, or by taking the individual electrode showing the strongest correlation.

## 3. Results

### 3.1 Performance on the continuous recall task

First, we assessed how increasing working memory load altered working memory performance. The Shapiro Wilk statistic confirmed the assumption of normality was violated for working memory capacity load 2 and all loads for precision, however, the sample size is robust to violations of normality (Tabachnick and Fidell 2018). Greenhouse Geiser statistics were used due to violation of sphericity. Working memory capacity was altered with load (*F*=74.0; *p*=1.2 x 10^-18^; ANOVA), with Bonferroni corrected post-hoc comparisons indicating that capacity increased from load 2 to load 4 (Δ*M* = −0.93; 95% CI =-1.07, −0.79; *t*(83) = −13.4; *p*=4.9 x 10^-22^, *d* = 1.46) but not from load 4 to load 6 (Δ*M* = −0.19; 95% CI =-0.39, 0.01; *t*(83) = −1.89, *p*=0.189, *d* = 0.21). Working memory precision was also altered with load (*F*=37.5, *p*=1.3 x 10^-9^; ANOVA), with standard deviations increasing (i.e., precision decreasing) from load 2 to load 4 (Δ*M* = −8.33; 95% CI = −9.77, −6.88; *t*(83) = −11.5; *p*=2.6 x 10^-18^, *d* = 1.25) and from load 4 to load 6, however this did not reach the threshold for significance (Δ*M* = −4.49; 95% CI = −8.23, −0.76; *t*(83) = −2.4, *p*=0.06, *d* = 0.26). Thus, our behavioural results are in line with other continuous recall and change detection tasks using coloured squares, showing a plateau in working memory capacity above 4 items (Luck and Vogel 2013).

### 3.2 Effect of task on aperiodic activity

Next, we assessed whether aperiodic activity changed during different periods of the working memory task. The aperiodic exponent differed from the baseline to the delay period in load 2 (*p =* 0.006, *d* = 0.51), load 4 (*p* = 0.002, *d* = 0.53) and load 6 (*p* = 0.003, *d* = 0.53), with exponent values increasing (i.e., slope steepening) in frontocentral electrodes, but decreasing (i.e., slope flattening) in lateral occipitoparietal electrodes (fig. 4). The aperiodic offset also decreased from the baseline to the delay period in occipitoparietal electrodes (fig. 4) in load 2 (*p* =0.005, *d* = −0.53), load 4 (*p* =0.007, *d* = −0.46), and load 6 (*p* =0.004, *d* =-0.50). Thus, both the aperiodic exponent and offset were altered during the delay period of working memory, however the direction of this change differed across the scalp.

**Figure 3.**
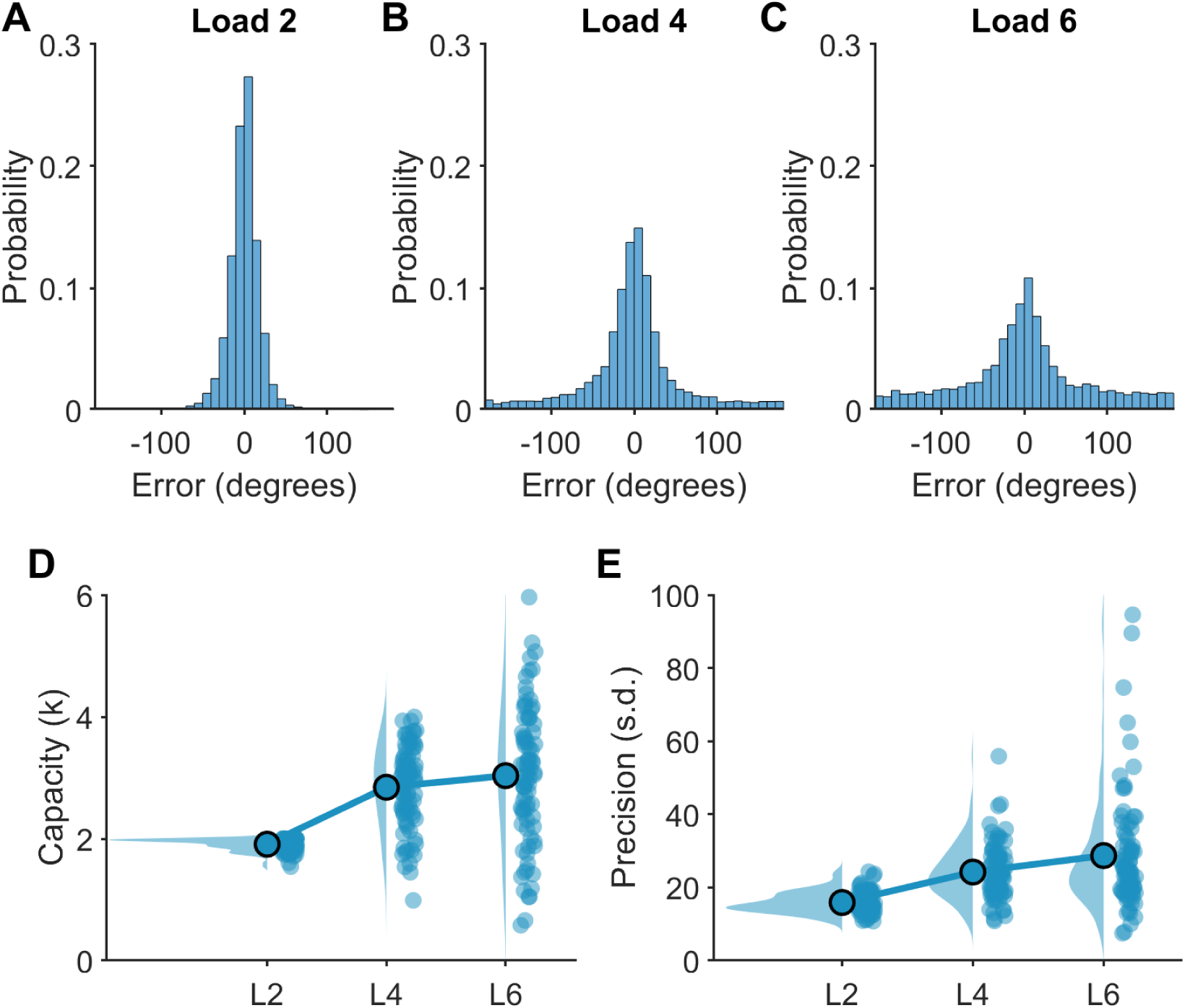
Working memory performance on the continuous recall task. A-C) Distribution of trial-by-trial performance calculated as error (distance from target in degrees) across all individuals (n=84) for each load. D-E) Working memory capacity (D) and precision (E) estimates at each load. Blue dots represent individual participants, highlighted blue dots indicate the group mean, and shaded blue area the distribution of the data.

**Figure 4.**
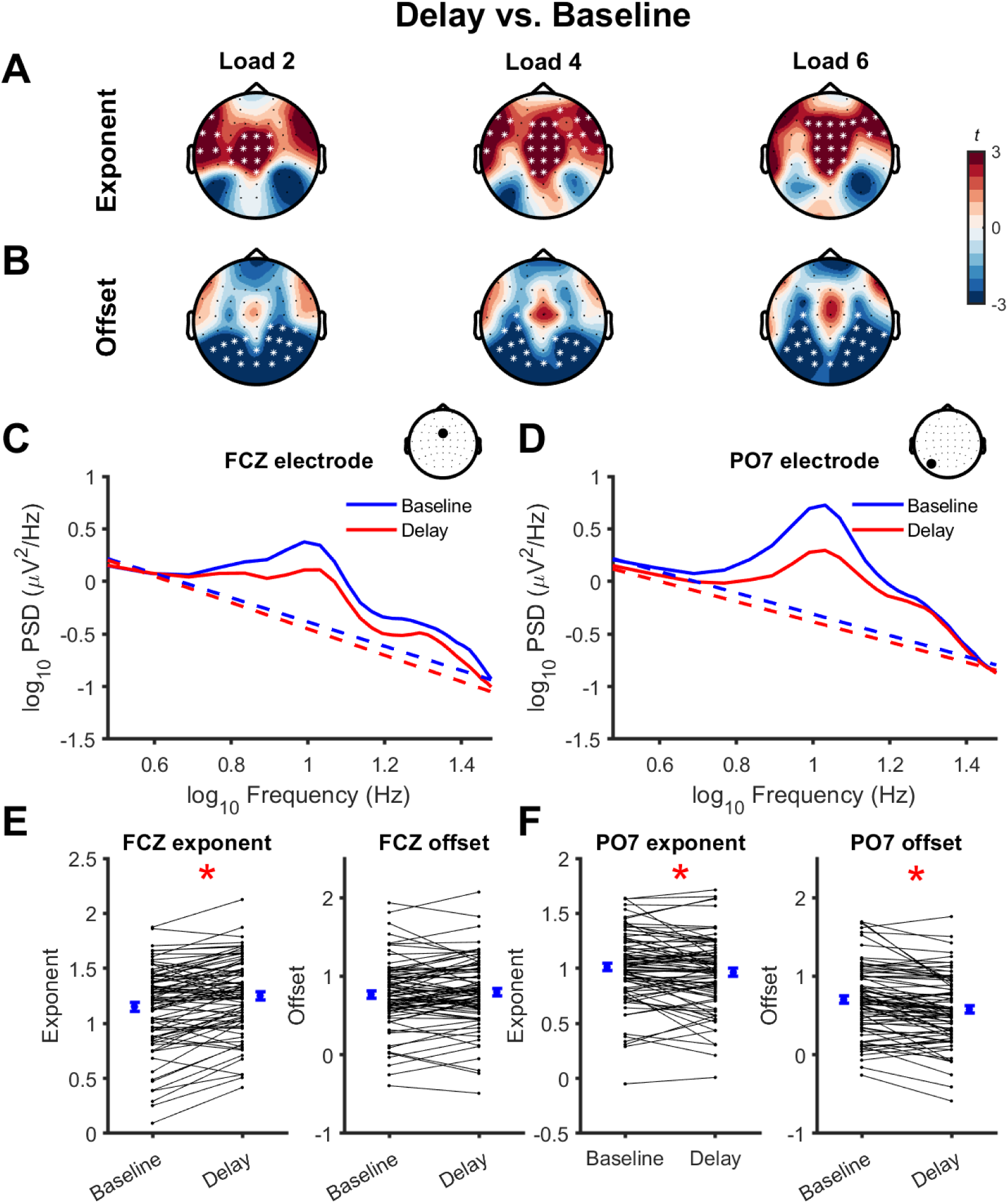
Task-related changes in aperiodic activity across the scalp. Topoplots representing changes (t-statistics) in aperiodic exponent (A) and offset (B) values between the baseline and delay periods of the working memory task at each load. * indicate electrodes contributing to significant clusters (*p*<0.01). C-D) Group mean EEG power spectra (solid lines) and aperiodic activity (dashed lines) from a frontocentral (C; FCZ) and lateral occipitoparietal (D; PO7) electrode during the baseline (blue) and delay (red) period of load 4. Data are plotted in log-log space. Inset topoplots indicate the position of the electrode on the scalp. E-F) Group mean (blue dots; ± standard error) and individual (black dots and lines) changes in aperiodic exponent and offset values from baseline to delay for frontocentral (E; FCZ) and lateral occipitoparietal (F; PO7) electrodes. * indicate a significant difference between time periods (*p*<0.05; paired t-test).

### 3.3 Effect of working memory load on aperiodic activity

Having established that aperiodic activity is altered during the delay period of working memory, we next asked whether the magnitude of these changes from the baseline to the delay period differed with increasing memory load. We could find no evidence for meaningful differences in task-related changes in aperiodic exponent or offset values when comparing memory loads (*no identified clusters*; fig. 5). These findings indicate that while the properties of aperiodic activity changed during the delay period, these task-related changes were of a comparable magnitude regardless of the memory load.

**Figure 5.**
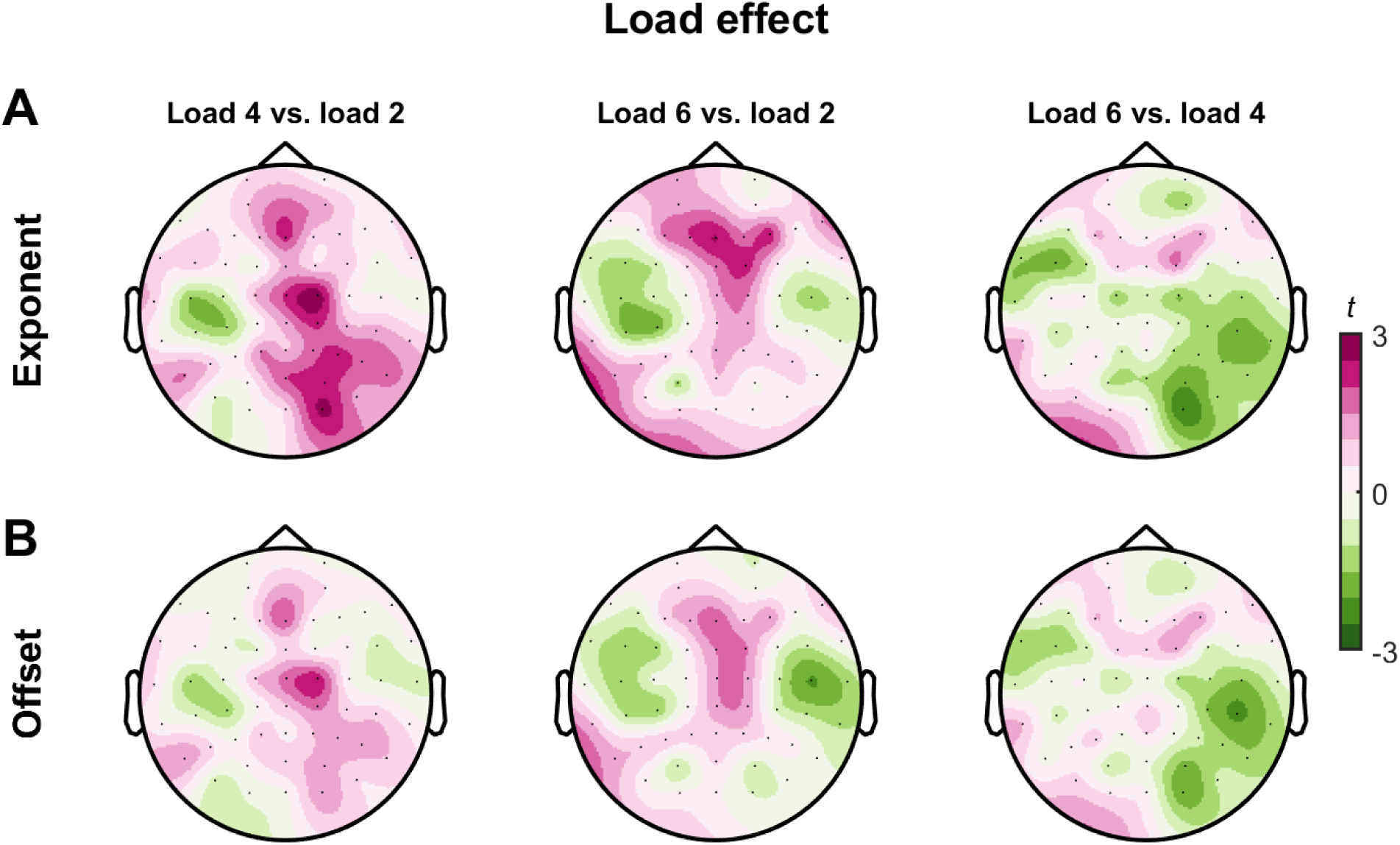
Load-dependent changes in task-related aperiodic activity. Topoplots representing differences (t-statistics) in task-related changes in aperiodic exponent (A) and offset (B) values between different memory loads. There were no significant clusters for any comparison (all *p*>0.05).

### 3.4 Relationship between changes in aperiodic activity and working memory performance

Next, we asked whether task-related changes from the baseline period to the delay period in aperiodic activity were related to working memory performance across individuals. Removal of outliers altered the sample size of analyses for the aperiodic exponent at load 2 (*n*= 83), load 4 (*n*=82), load 6 (*n*=80), and the set size effect (*n*=82) and for the offset at load 2 (*n*=81), load 4 (*n*=83) and load 6 (*n*=81) and the set size effect (*n*=83). Task-related changes in aperiodic exponent values were positively associated with working memory capacity at load 2 (*p* = 0.039, *r* = 0.31), load 4 (*p* = 0.015, *r* = 0.26) and load 6 (*p* = 0.024, *r* = 0.25), such that changes towards steeper aperiodic slopes were associated with higher working memory capacity. However, this did not reach the adjusted threshold for significance at load 2 or 6 (fig. 6A-F). Although we didn’t observe any load-dependent changes in aperiodic activity (fig. 5), we also assessed whether set-size effects (the mean change in aperiodic slope for supra-capacity loads 4 and 6 subtracted from sub-capacity load 2) were related to working memory capacity (quantified as *Kcd*; the mean capacity for suprathreshold loads 4 and 6), as these relationships have been reported for other EEG measures (Fukuda, Mance, and Vogel 2015). We could not find evidence for a relationship between working memory capacity and the set size effect (*no significant clusters*; fig. 6G-H). In contrast to slope, we could not find evidence for meaningful associations between aperiodic offset and capacity at any load (load 2, *p* = 0.061; load 4, *p*= 0.115; load 6, *p*= 0.196, set-size effect, *no clusters*), or between any of the parameters and working memory precision (*no significant clusters*). These findings indicate that individuals who showed larger increases from the baseline period to the delay period in aperiodic exponent (i.e., steeper slopes) in frontocentral electrodes at supra-capacity loads have a higher working memory capacity.

**Figure 6.**
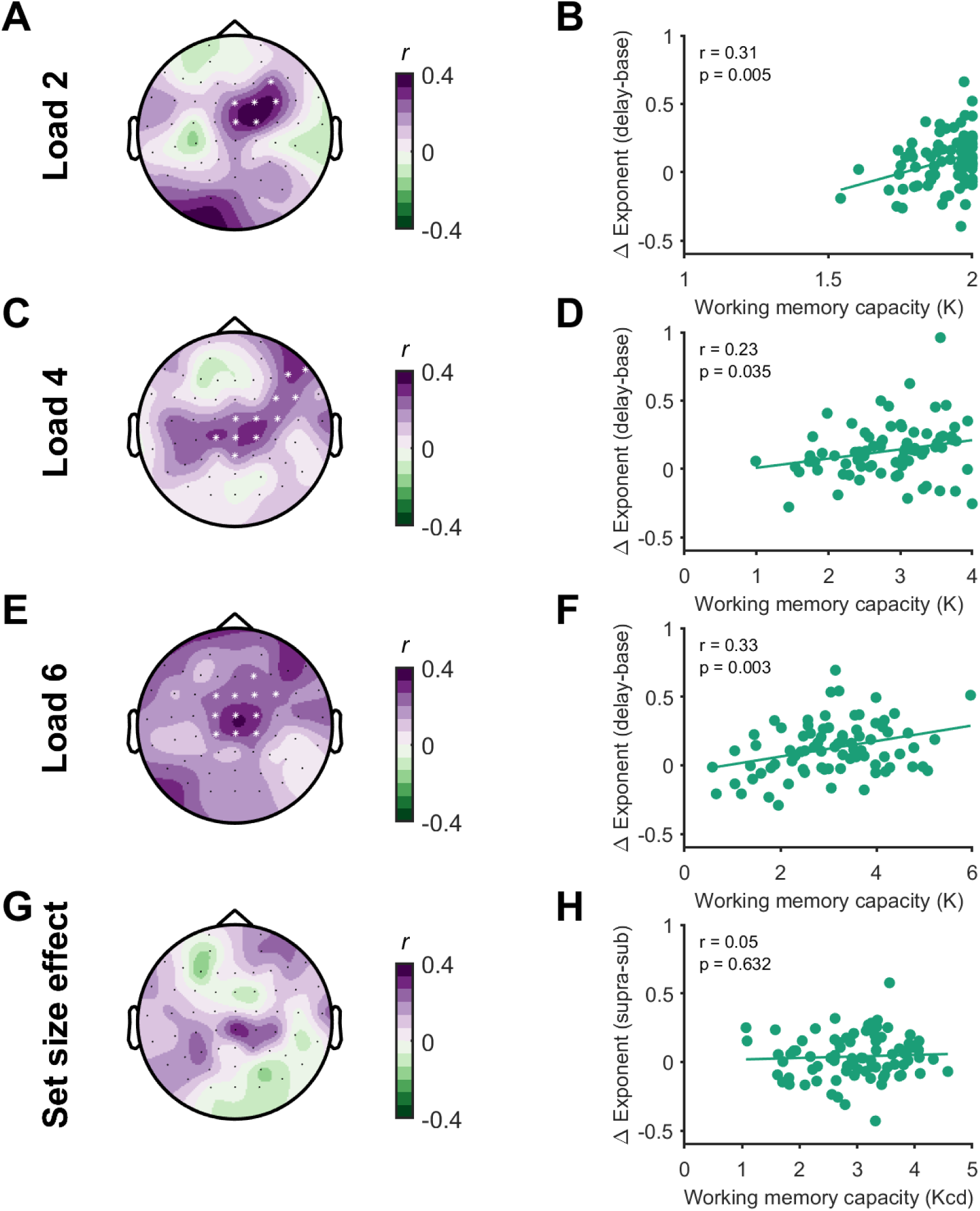
Relationship between task-related changes in aperiodic exponent and working memory capacity. Topoplots demonstrating the relationship (Pearson’s correlation, r) between task-related changes in the aperiodic exponent and working memory capacity at each electrode (left column), and scatter plots for a central channel (Cz, right column). Data are plotted for each load and the set size effect. * indicate channels contributing to significant clusters (*p*<0.05).

We then asked whether the relationship between changes in aperiodic activity and capacity generalised to measures of working memory ability from complex span tasks. Factor score estimates of complex span performance were related to suprathreshold working memory capacity (Kcd) from the continuous recall task (*p* = 0.018; *r* = 0.27) and individual capacity estimates from load 2 (*p* = 3.75×10^-4^, *r* = 0.40), load 4 (*p* = 0.001, *r* = 0.37), but not load 6 (*p* = 0.137, *r* = 0.17), indicating performance on the colour wheel task is related to broader measures of working memory capacity. Additionally, estimates of complex span performance were also positively associated with task-related changes in aperiodic exponent at load 4 (*p* = 0.015, one-tailed; *r* = 0.24), but not load 2 (*p* = 0.170), load 6 (*p* = 0.06) or for the set-size effect (*no significant clusters*) (fig. 7). Together, these findings suggest that task-related changes in aperiodic activity index a neural mechanism which is important for general working memory ability, as opposed to simply indexing task-specific effects.

**Figure 7.**
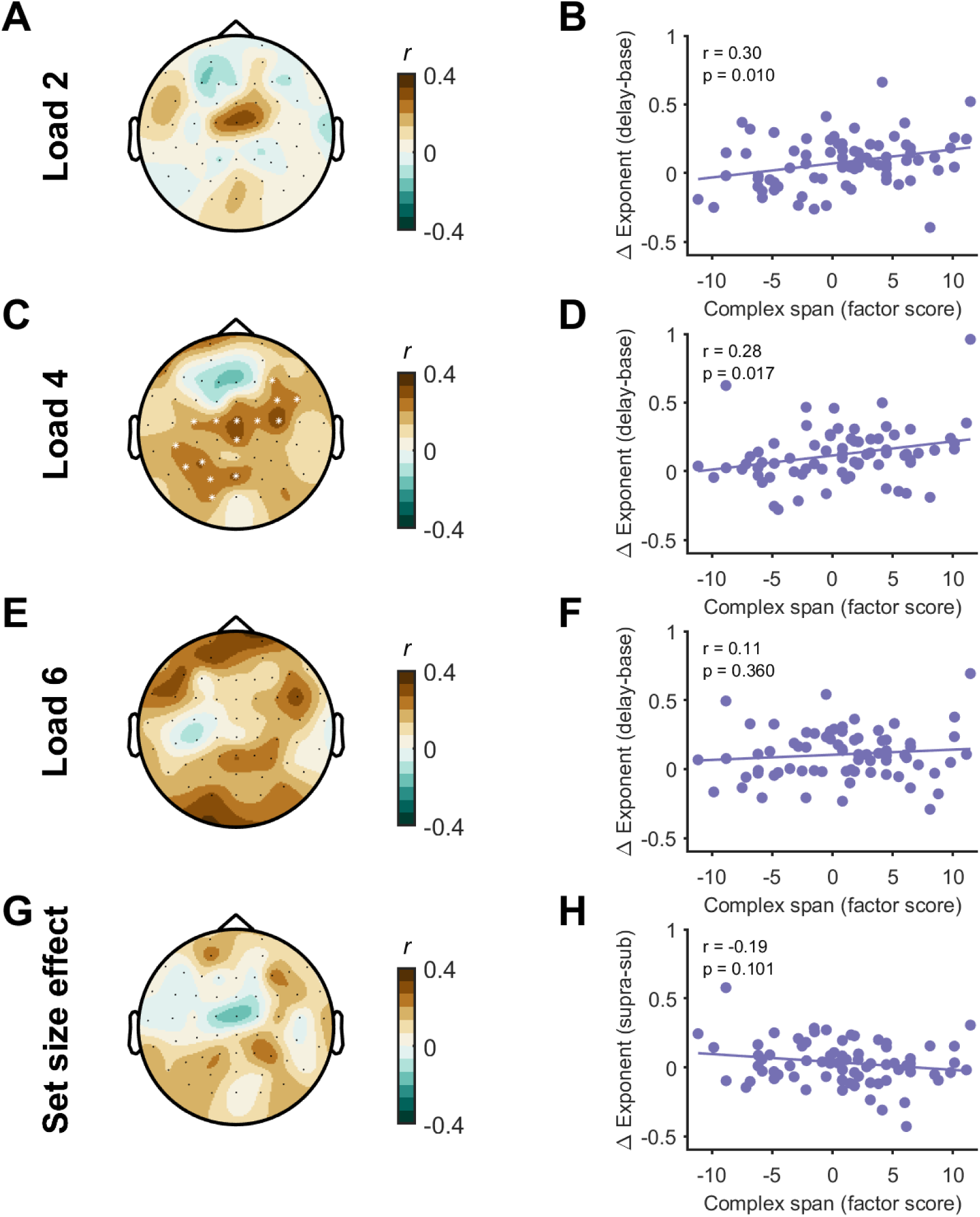
Relationship between task-related changes in the aperiodic exponent and complex span working memory capacity. Topoplots demonstrating the relationship (Spearman’s rho, r_s_) between task-related changes in the aperiodic exponent and complex span working memory capacity (represented as a factor score) at each channel (left column), and scatter plots from a central channel (Cz; right column). Data are plotted for each load and the set size effect. * indicate channels contributing to significant clusters (*p*<0.05).

### 3.5 Alpha-band oscillatory power and slow wave ERPs

Next, we attempted to replicate previously established EEG correlates of working memory capacity (Fukuda, Mance, and Vogel 2015), including alpha-band oscillatory power and slow-wave ERPs. Cluster-based permutation analyses showed that alpha-band oscillatory power was reduced from the baseline to the delay period across all electrodes for each load (load 2, *p* =3.99 x10^-4^, *d* = 0.57; load 4, *p* =3.99 x10^-4^, *d* = 0.52; load 6, *p* =3.99 x10^-4^, *d* = 0.60). This suppression was stronger for load 4 compared to load 2 with electrodes included in the significant cluster found across right hemisphere and centro-parietal electrodes (*p* = 0.044, *d* = 0.32), but did not differ between load 4 and load 6 (*no identified clusters*; fig. 8). Removal of outliers altered the sample size of correlation between alpha suppression and working memory capacity for the set size effect (*n*=79). The alpha suppression set-size effect had a negative association with working memory capacity (*p* = 0.022, *r* = −0.27; 8E,G), indicating that stronger alpha suppression at supra-capacity loads was related to higher capacity. We did not find any associations between alpha suppression and complex span working memory capacity (n=71, *p =* 0.109, fig. 8F,H), although there was a weak relationship across similar channels. Together, these findings replicated well-established capacity-limited effects on alpha suppression and relationships between alpha suppression and working memory capacity (Fukuda, Mance, and Vogel 2015).

**Figure 8.**
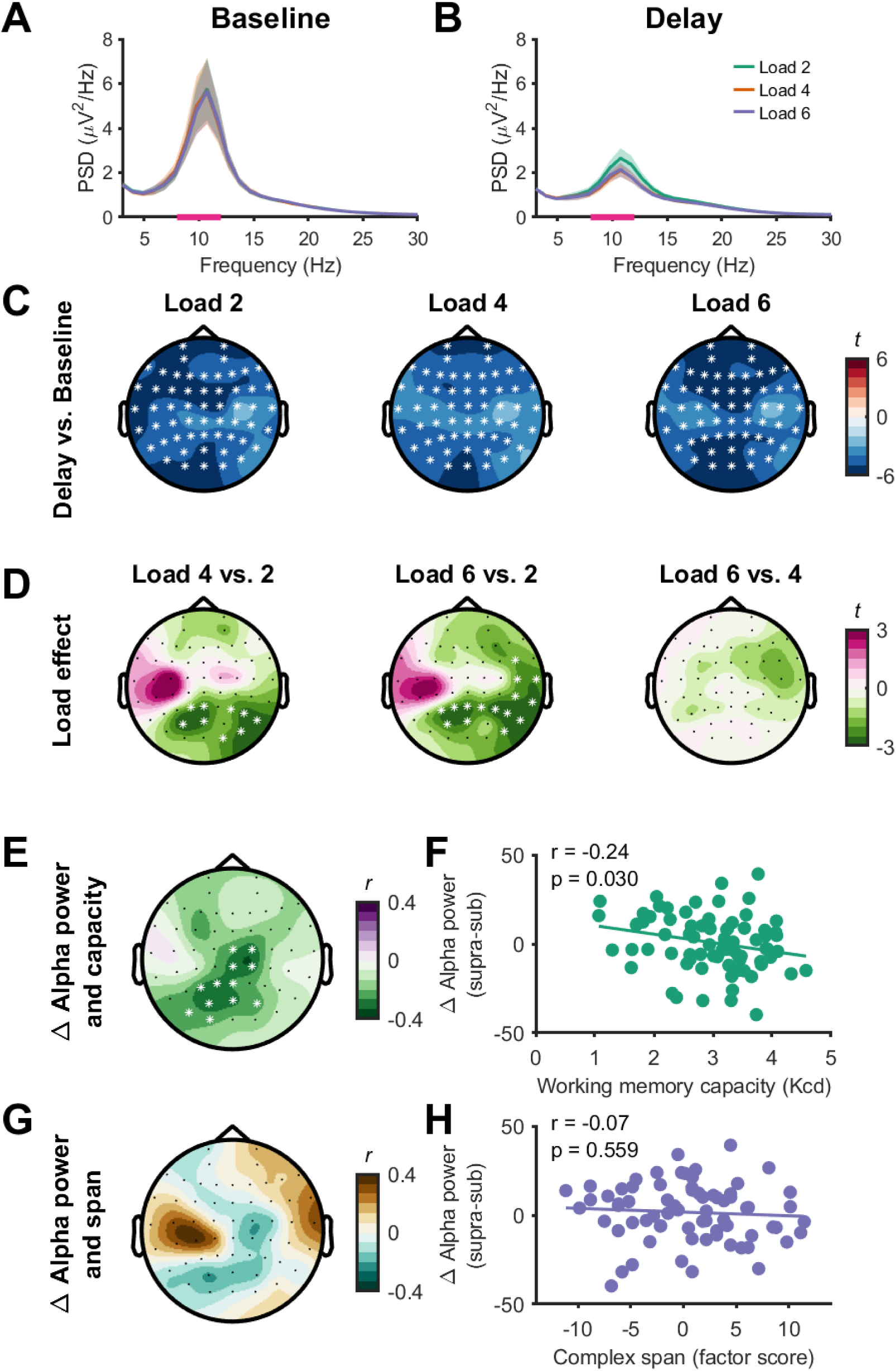
Changes in alpha-band oscillatory power with working memory. A-B) EEG power spectra (power spectral density, PSD) during the baseline (A) and delay (B) period for each load (channel P8). Solid line represents the group mean and shaded bars the standard error. Pink bars on the x axis show the frequency range of interest. C) Topoplots indicating changes (t-statistics) in alpha-band oscillatory power between the baseline and delay periods across all electrodes. D) Topoplots indicating differences (t-statistics) in task-related alpha suppression between loads across all electrodes. E-H) Topoplots demonstrating the relationship (Pearson’s correlation, r) between task-related changes in alpha-oscillatory power and working memory capacity (E) and complex span performance (G) at each electrode (left column), and scatter plots for a central electrode (Cz, right column; F and H). Data are plotted for the set size effect. * indicate electrodes contributing to significant clusters (p<0.05).

Slow wave ERPs during the delay period increased in amplitude between load 2 to load 4 with the significant cluster involving frontal electrodes, (*p* = 0.032 *d* = 0.39), and decreased in amplitude in another cluster that involved parietal electrodes (*p* = 3.999 x10^-4^, *d* = −0.55, fig. 9). However, there was no significant difference in the slow wave ERP between load 4 to load 6 (*p* = 0.136), indicating a plateau in slow-wave ERP amplitude changes at supra-capacity loads. Removal of outliers altered the sample size of correlations between the slow wave ERP and working memory set size effect (*n*=79). The slow wave ERP set-size effect did not show a significant relationship with working memory capacity at the cluster level (*p =* 0.173), although qualitatively a greater reduction in ERP amplitude was negatively associated with higher working memory capacity in some frontal electrodes (fig. 9E-F). Furthermore, the slow wave ERP set size effect (*n*=71) was associated with complex span performance (*p* = 0.010, *r =* −0.30) showing a negative relationship between slow wave ERP amplitude change and performance over frontal electrodes and a positive relationship over occipital electrodes (fig. 9G-H). Together, these findings replicate established capacity-limited load effects on slow wave ERP amplitudes and relationships between changes in slow-wave ERPs with loads and broader measures of working memory capacity (Fukuda, Mance, and Vogel 2015; Unsworth et al. 2015).

**Figure 9.**
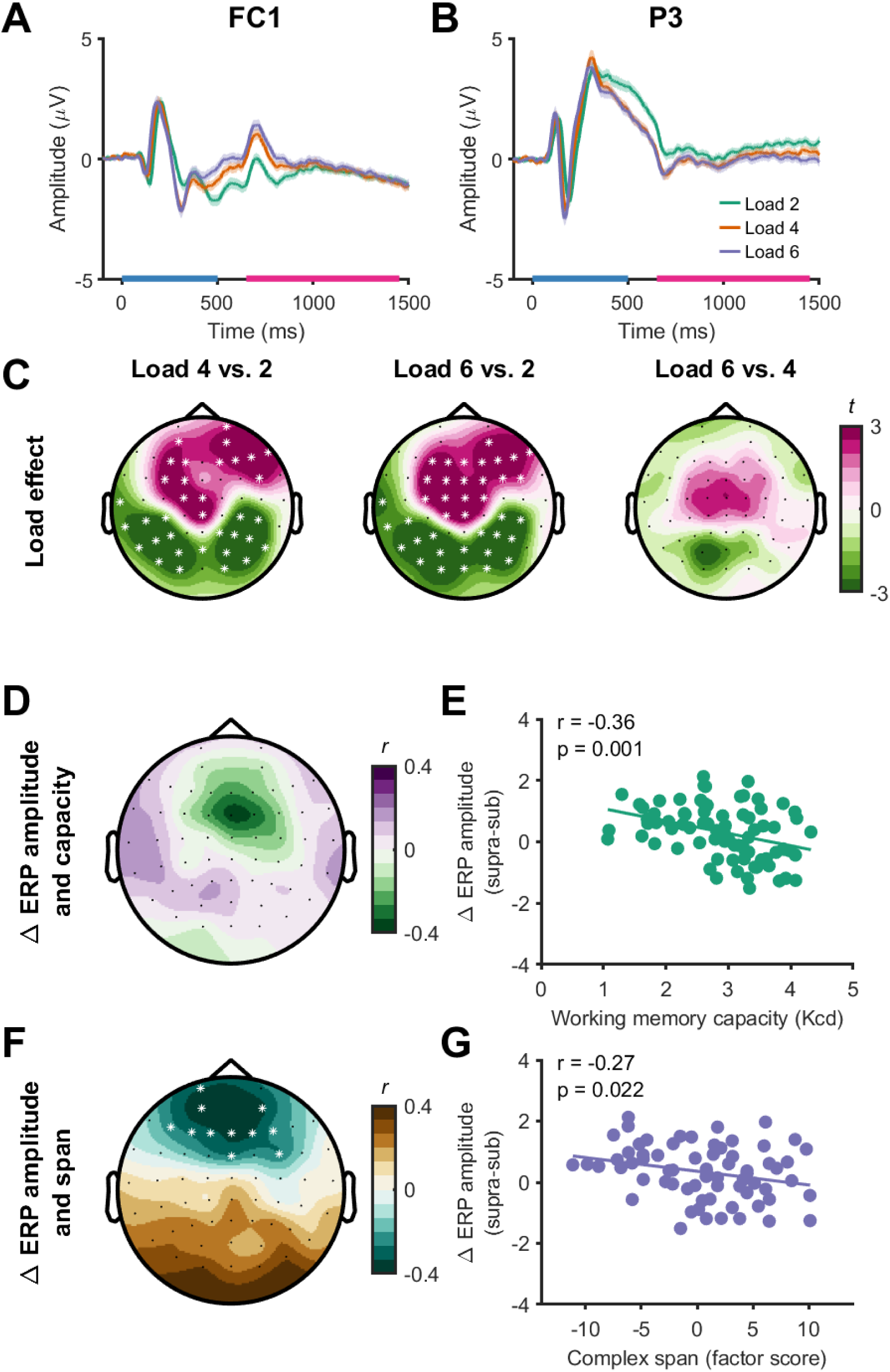
Changes in slow-wave event-related potentials (ERPs) with working memory. A-B) ERPs from a left frontal (FC1, A) and parietal (P3, B) electrode at different working memory loads. The blue bar on the x-axis represents the timing of the stimulus (0-500 ms), with the slow-wave ERP period being analysed from 650-1450 ms (pink bar on the x-axis). C) Topoplots indicating differences in slow-wave ERP amplitude during the delay period between loads across all electrodes. D-G) Topoplots demonstrating the relationship (Pearson’s correlation, r) between slow-wave ERP amplitude changes with load and working memory capacity (D) and complex span performance (F) at each electrode (left column), and scatter plots for a frontocentral electrode (FCz, E and G). Data are plotted for the set-size effect. * indicate electrodes contributing to significant clusters (*p*<0.05).

### 3.6 Relationship between EEG measures and working memory capacity

Finally, we asked whether the different EEG measures explained independent variance in working memory capacity. To examine the independent variance in working memory capacity explained by each EEG measure, we entered either: 1) the average of all electrodes that contributed to a significant cluster; or 2) the electrode showing the peak correlation with working memory capacity as variables into a multiple linear regression model that tested the relationship between neural activity set size effects and working memory capacity (*Kcd*). Using the average of electrodes in a cluster explained 21.5% of variance (*p* = 2.08 x10^-4^; n = 72) in working memory capacity, with all variables providing independent contributions to the relationship. Similarly, using the peak correlation electrode explained 23.3% of variance (*p* = 9.78 x10^-5^; n = 72) with all variables providing independent contributions to the relationship (table 1). For complex span performance, using the average of electrodes in a cluster explained 27.1% of variance (*p* = 6.27 x10^-5^; n = 64) with the aperiodic exponent and slow-wave ERP providing independent contributions to the relationship. Using the peak correlation electrode also explained 27.1% of variance (*p* = 6.23 x10^-5^; n = 64) in complex span performance again with the aperiodic exponent and slow-wave ERP providing independent contributions to the relationship (table 2). Together, these findings suggest that changes in the aperiodic exponent, slow-wave ERPs and, to a lesser extent, alpha suppression index neural mechanisms that make distinct contributions to individual differences in working memory capacity measured across different tasks.

**Table 1.** Parameter estimates from the multiple linear regression model for working memory capacity as a function of EEG measures.

| Variables | *Average cluster* |  | *Peak correlation* |  |
| --- | --- | --- | --- | --- |
| | $\beta$ | $p$ | $\beta$ | $p$ |
| Intercept | 2.86 | $5.7 \times 10^{-39}$ | 2.90 | $1.6 \times 10^{-42}$ |
| Aperiodic exponent | 1.41 | 0.05 | 1.43 | $9.1 \times 10^{-3}$ |
| Alpha suppression | - 0.02 | 0.01 | - 0.02 | 0.01 |
| Slow-wave ERP | - 0.25 | 0.03 | - 0.23 | 0.02 |

**Table 2.** Parameter estimates from the multiple linear regression model for complex span performance as a function of EEG measures.

| Variables | *Average cluster* |  | *Peak correlation* |  |
| --- | --- | --- | --- | --- |
| | $\beta$ | $p$ | $\beta$ | $p$ |
| Intercept | 0.26 | 0.68 | 0.48 | 0.42 |
| Aperiodic exponent | 10.96 | 0.03 | 1.43 | $3.1 \times 10^{-3}$ |
| Alpha suppression | 0.05 | 0.14 | - 0.02 | 0.58 |
| Slow-wave ERP | - 0.26 | $8.6 \times 10^{-4}$ | - 0.23 | $4.3 \times 10^{-4}$ |

## 4. Discussion

The aim of this study was to comprehensively assess changes in broadband aperiodic activity in the EEG power spectrum during a visual working memory task. We found that characteristics of aperiodic activity were altered during the delay period of the task compared to a baseline period, with the aperiodic exponent increasing at fronto-central electrodes (i.e., slope of the power spectrum steepening), and decreasing (i.e., slope flattening) at lateral parieto-occipital electrodes. The broadband offset of the power spectrum also decreased across parieto-occipital electrodes during the delay period. Importantly, the task-related changes in aperiodic activity were not modulated by different working memory loads. Across individuals, a larger task-related increase in the aperiodic exponent (i.e., steeper slope) over fronto-central electrode during supra-capacity loads (4 or 6 items) was associated with higher working memory capacity, a relationship which extended to performance in complex span working memory tasks collected separately from the EEG task. Finally, the relationship between task-related changes in the aperiodic exponent and working memory capacity was independent of other EEG correlates of capacity, including the suppression of alpha-band oscillatory activity and slow-wave ERPs. Together, these findings suggest that task-related changes in the aperiodic exponent reflect a dissociable neural mechanism that is important for working memory ability.

The broadband 1/f-like power distribution is a well-recognized characteristic in neural recordings and is observed across numerous frequency ranges and different data types, including local field potentials, intracranial and scalp EEG, magnetoencephalography, and functional MRI (He 2014). However, the physiological relevance of this broadband aperiodic activity has been often overlooked in previous literature, with most analysis methods measuring narrow-band frequency ranges which potentially conflate rhythmic periodic activity with aperiodic activity (Donoghue et al. 2020). Studies which separate broadband aperiodic activity from periodic activity have demonstrated that aperiodic features of neural recordings are altered across a range of short and long timescales including in response to sensory stimuli and during motor responses (Miller et al. 2014; Podvalny et al. 2015; Waschke et al. 2021), as well as from rest to task engagement (Donoghue et al. 2020), from wake to sleep (Lendner et al. 2020), and across the human lifespan (Schaworonkow and Voytek 2021; Voytek et al. 2015; Merkin et al. 2023). For example, Podvalny and colleagues (2015) found a reduction in the aperiodic exponent (i.e., flattening of the power spectrum slope) in intracranial EEG recordings taken from task-related brain regions following visual, auditory, and motor events. Similar to our results, Donoghue and colleagues (2020) found a reduction in both the aperiodic exponent and offset in scalp EEG recordings over lateral parieto-occipital regions during the delay period of a visual working memory task. We have expanded upon these findings by demonstrating that the changes in the aperiodic exponent during the delay period of a working memory task are dependent on the scalp measurement location, with the aperiodic exponent increasing at fronto-central electrodes. Interestingly, the task-related changes in aperiodic exponent and offset were not modulated by different working memory loads, suggesting the differences in aperiodic activity may reflect a general change in task-related brain state which is independent of the number of items held in memory. Gyurkovics and colleagues (2022) found that the slope of the EEG power spectra became steeper following novel as opposed to repeated auditory stimuli during an oddball task (Gyurkovics et al. 2022). As such, changes in the slope of the spectra may reflect a neural mechanism sensitive to attentional demands. Together, our findings support a growing body of evidence that aperiodic activity in the EEG power spectrum is modulated by cognitive tasks, possibly reflecting a mechanism reflecting attention. Furthermore, we’ve added to this literature by showing that the direction of changes in the aperiodic slope differs between posterior and frontal regions.

Task-related changes in the slope of the power spectrum seem to reflect a physiological mechanism important for working memory. Several studies have reported a relationship between the aperiodic exponent and working memory performance, including from both scalp and intracranial EEG recordings (Gao et al. 2020; Donoghue et al. 2020; Voytek et al. 2015; Lendner et al. 2023; Sheehan et al. 2018). Importantly, Donoghue and colleagues (2020) showed that a model that included task-related changes in aperiodic exponent and offset values was better able to explain working memory performance compared to models that included other neural correlates of working memory such as changes in alpha-band oscillatory power. We partially replicated and extended this finding, showing that larger task-related increases in aperiodic exponent values were related with higher working memory capacity independent of both alpha power suppression and changes in slow-wave ERP amplitude. Alpha power suppression and lateralized slow-wave ERPs (known as contralateral delay activity) were once thought to measure a common neural mechanism (van Dijk et al. 2010). However, more recent research suggests the measures are not related to one another in non-lateralised tasks, exist over a different time course, and explain independent variance in working memory capacity across individuals (Fukuda, Mance, and Vogel 2015). In addition, studies using multivariate pattern analysis have shown that both stimulus position and feature information can be decoded from slow-wave ERPs, but only stimulus position from alpha-band oscillatory power (Bae and Luck 2018; Foster et al. 2016; 2017), suggesting the measures reflect different neural mechanisms relating to different aspects of working memory. Beyond task-specific performance, lateralised slow-wave ERPs during the delay period are related to measures of fluid intelligence, long-term memory, and attention control (Unsworth et al. 2015), suggesting this mechanism is important for broader cognitive ability. We have shown that the relationship between increases in the aperiodic exponent and capacity also extends beyond the task used in the EEG experiment, relating to measures of working memory performance obtained from complex span tasks. Complex span and non-selective visual array tasks like the continuous recall task in the current EEG experiment (i.e., tasks that do not contain any distractor stimuli) share common variance which is thought to primarily reflect working memory storage capacity, with a small contribution from attention control (Draheim et al. 2021; Martin et al. 2021; Shipstead, Harrison, and Engle 2015). Thus, our findings suggest that changes in the aperiodic exponent reflect a neural mechanism distinct from alpha power suppression and slow-wave ERPs which may be important for determining storage capacity limits in working memory. Future work measuring a broad range of cognitive tasks is required to further clarify the relationship between changes in the aperiodic exponent and other aspects of cognition relevant for working memory like attention.

What are the neural mechanisms that could contribute to changes in aperiodic activity measured in the EEG power spectrum during visual working memory? A number of neurophysiological properties have been suggested to underlie 1/f-like activity observed in EEG, including the dynamic balance of excitation and inhibition (Gao, Peterson, and Voytek 2017), low-pass filtering properties of dendrites on cortical excitatory neurons (Lindén, Pettersen, and Einevoll 2010) or the extracellular volume (Bédard, Kröger, and Destexhe 2006), the connectivity structure of neural networks (Chaudhuri, He, and Wang 2018), or the sum of stochastically-driven damped linear oscillators tuned to different alpha band frequencies (S. D. Muthukumaraswamy and Liley 2018; Evertz, Hicks, and Liley 2022). Gao et al. (2017) used a biophysical model of excitatory and inhibitory neural populations and physiological data to demonstrate that the slope of the power spectrum measured by the aperiodic exponent in invasive recordings is sensitive to changes in the balance of excitatory and inhibitory neurotransmission, with a flattening slope indicating a greater excitation/inhibition balance and vice versa. In non-invasive recordings including EEG and MEG, anaesthetics and other pharmacological agents targeting excitatory and inhibitory receptors also alter the aperiodic exponent (Colombo et al. 2019; S. D. Muthukumaraswamy and Liley 2018), providing further evidence that the excitation/inhibition balance can be assessed by aperiodic slope detected in scalp signals, although not always in the expected direction (Salvatore et al. 2024; Brake et al. 2024). Thus, one possible interpretation of changes in the aperiodic exponent observed in our study is a shift towards excitation over parieto-occipital regions and inhibition over frontal brain regions. Indeed, several recent studies using optogenetics (D. Kim et al. 2016), and recurrent neural networks (R. Kim and Sejnowski 2021) have suggested that increased inhibitory-inhibitory neurotransmission in frontal regions could underlie a lengthening of neural timescales which stabilises working memory performance. Regardless of the physiological mechanism, a shift towards slower frequencies results in long term autocorrelations, which may represent a putative physiological mechanism capable of carrying memory information during visual working memory tasks (Wasmuht et al. 2018; Gao et al. 2020).

In contrast, changes in the broadband offset of the power spectrum in invasive recordings have been linked with changes in neuronal firing rates (Manning et al. 2009), with increased firing rates associated with increased offsets and vice versa. Thus, the reduction in the broadband offset over posterior regions may reflect a reduction in neuronal firing rates in parieto-occipital regions during the working memory delay period. Invasive recordings from visual (van Kerkoerle, Self, and Roelfsema 2017) and parietal (Pesaran et al. 2002) regions in non-human primates typically suggest that neuronal firing rates either increase in task-relevant neurons, or return to baseline levels during the delay period, inconsistent with the above interpretation of our findings. The relationship between local-field potentials and scalp EEG can be complex (Snyder, Issar, and Smith 2018) and simultaneous invasive and non-invasive recordings are required to further clarify how changes in neural firing rates impact EEG power spectra measured at the scalp.

There are several limitation to our study. First, the EEG power spectra is contaminated by scalp muscle activity (S. Muthukumaraswamy 2013), which could impact measures of aperiodic activity including the aperiodic exponent and offset (Gerster et al. 2022). We attempted to minimise this contamination by using a validated method to select and remove independent components likely representing muscle activity (Fitzgibbon et al. 2016), and by limiting our quantification of the power spectrum to lower frequencies (<30 Hz) which are thought to be less effected by muscle activity. However, these approaches are not foolproof, and also limit our capacity to detect changes in the higher-frequency ranges of the power spectrum, which likely carry important information on neural changes during working memory. Second, it is unclear from our study whether the changes in aperiodic activity are specific to working memory processes or reflect general changes in brain activity related to sensory processing or attention. Indeed, the lack of modulation of aperiodic activity with increasing memory load may indicate that these changes are not specific to working memory and could reflect other cognitive processes like attention (Waschke et al. 2021). Nonetheless, changes in the aperiodic exponent were related to performance across tasks, suggesting an important role for this neural mechanism in general working memory ability regardless of the specific sensory or cognitive process it underlies. Third, we used one specific model (SpecParam/FOOOF) to quantify the aperiodic features of the power spectra. While this method is widely used and has been extensively validated using simulated data (Donoghue et al. 2020), studies have demonstrated that the aperiodic slope fits can overestimates power at certain frequencies, resulting in oscillatory power estimates being negative – a physical impossibility (see fig. 9A-B for examples) (Barry and Blasio 2021). Several different modelling approaches exist for quantifying the aperiodic 1/f-like characteristics of EEG data including IRASA (Wen and Liu 2016) and PaWNextra (Barry and Blasio 2021), which have differing strength and weaknesses relative to SpecParam/FOOOF. Future work would benefit from cross referencing these different methods for quantifying changes in the aperiodic features of EEG power spectra during cognitive tasks. Finally, we used set time periods during the baseline and the delay periods of the task to quantify the EEG power spectra and estimate aperiodic features of the signal. More recently, algorithms have been developed which allow measurement of periodic and aperiodic features across time (e.g., SPRiNT, Spectral Parameterization Resolved in Time) (Wilson, da Silva Castanheira, and Baillet 2022). Using such methods would allow for a more fine-grained estimation of changes in aperiodic activity across time during working memory and other cognitive tasks.

Overall, we provide evidence for region-specific changes in aperiodic activity during the delay period of a visual working memory task. Task-related changes in the slope of the power spectrum measured by the aperiodic exponent likely reflect a neural mechanism distinct from oscillatory and ERP changes also observed during the task and appear important for general working memory ability. Our findings add to a growing body of evidence demonstrating the physiological relevance of aperiodic activity in brain function and underscore the importance of separating periodic and aperiodic activity when quantifying changes in EEG power spectra during cognitive tasks.

## 5.1 Data and code availability

Data used for the statistical analyses and to generate figures are available from: https://figshare.com/articles/dataset/Aperiodic_activity_during_working_memory/29146721 Code used for analyses is available from: https://github.com/nigelrogasch/aperiodic-activity-working-memory

## 5.2 Author contributions

**Sian Virtue-Griffiths:** Conceputalisation, Software, Formal analysis, Investigation, Data Curation, Writing – Original Draft, Visualisation. **Alex Fornito:** Conceputalisation, Methodology, Writing - Review & Editing, Supervision, Project Administration, Funding Acquisition. **Sarah Thompson**: Investigation, Data Curation, Writing - Review & Editing. **Mana Biabani:** Investigation, Data Curation, Writing - Review & Editing. **Jeggan Tiego:** Formal Analysis, Writing - Review & Editing. **Tribikram Thapa:** Formal Analysis, Writing - Review & Editing. **Neil W. Bailey:** Writing - Review & Editing. **Nigel C. Rogasch:** Conceputalisation, Methodology, Software, Validation, Formal analysis, Investigation, Data Curation, Writing – Original Draft, Visualisation, Supervision, Project Administration, Funding Acquisition.

